# The Rab32/BLOC-3 dependent pathway mediates host-defence against different pathogens in human macrophages

**DOI:** 10.1101/570531

**Authors:** Massimiliano Baldassarre, Virtu Solano-Collado, Arda Balci, Rosa A. Colamarino, Ivy M Dambuza, Delyth M. Reid, Heather M Wilson, Gordon D Brown, Subhankar Mukhopadhyay, Gordon Dougan, Stefania Spanò

**Author notes:** the authors equally contributed to the experimental part of this paper (Massimiliano Baldassarre, Virtu Solano-Collado).

## Abstract

Macrophages provide a first line of defence against microorganisms, and while some mechanisms to kill pathogens such as the oxidative burst are well described, others are still undefined or unknown. Here we report that the Rab32 GTPase and its guanine nucleotide exchange factor BLOC-3 are central components of a trafficking pathway that controls both bacterial and fungal intracellular pathogens. This broad host-defence mechanism is active in both human and murine macrophages and is independent of well known antimicrobial mechanisms such as the NADPH-dependent oxidative burst, production of nitric oxide and antimicrobial peptides. To survive in human macrophages, *Salmonella* Typhi actively counteracts the Rab32/BLOC-3 pathway through its *Salmonella* pathogenicity island-1-encoded type III secretion system. These findings demonstrate that the Rab32/BLOC-3 pathway is a novel and universal host-defence pathway and protects mammalian species from a wide range of intracellular pathogens.

## INTRODUCTION AND RESULTS

Cells of our innate immune system e.g. macrophages are involved in the first line of defense against microorganisms. After phagocytosis, macrophages can eliminate most of the microorganisms they encounter by directing them in intracellular compartments where conditions are not compatible with microorganism life. A key strategy used by macrophages to kill microorganisms is the production of reactive oxygen species (ROS) through activation of the NADPH oxidase complex that is assembled on cellular membranes in response to infection (Panday et al., 2015). Other mechanisms, such as the production of nitric oxide, or cathelicidin-related antimicrobial peptides can also mediate bacterial killing (Fang 2004; Xhindoli et al. 2016). Despite the presence of a number of effective antimicrobial mechanisms, some microorganisms have evolved to become effective intracellular pathogens by escaping clearance and killing mechanisms present in macrophages and other immune cell types. For example, *Salmonella enterica* harbours two type III secretion systems that are responsible for the delivery of a battery of effectors that allow *Salmonella* to invade host cells, including macrophages, and survive in a specialized intracellular compartment known as the *Salmonella*-containing vacuole (SCV) (Galan et al., 2014; Jennings et al., 2017; Hume et al., 2017). This broad bacterial species is genetically diverse and includes hundreds of different serovars that can cause human and important veterinary diseases. *Salmonella enterica* serovar Typhi (*S*. Typhi) is a human-restricted serovar that causes typhoid fever, a disease that affect ≈22 million people every year (Waddington et al., 2014). Unlike many *Salmonella* serovars that can infect a broad-range of hosts, *S*. Typhi naturally only infects humans (Dougan and Baker, 2014). For example, it cannot establish an oral infection in laboratory mice (Spanò and Galan, 2012).

Previously, we have shown that the inability of *S*. Typhi to infect mice depends, at least in part, on the fact that this pathogen cannot target the Rab32 GTPase (Spanò, 2016) in mouse macrophages. This GTPase and its guanine nucleotide exchange factor BLOC-3 (<u>B</u>iogenesis of <u>L</u>ysosome-related <u>O</u>rganelles <u>C</u>omplex-3) are central components of a pathway that regulate membrane trafficking to lysosome-related organelles in several specialized cell types (Spanò, 2016; Spanò and Galan, 2012). The murine Rab32/BLOC-3 pathway is effectively neutralized by the murine-virulent *Salmonella enterica* serovar Typhimurium (*S*. Typhimurium) through the delivery of two SPI-2 type III secretion effectors, GtgE and SopD2, that directly target Rab32 by acting as a protease and a GTPase activating protein, respectively(Spanò and Galan, 2012; Spanò et al., 2016). A *S*. Typhimurium mutant defective for both these effectors is virtually avirulent in wild type mice, but is able to infect mice that are either deficient for Rab32 or BLOC-3 (Spanò et al., 2016).

This suggested that the Rab32/BLOC-3 pathway limits the infectivity of bacteria that have not evolved to neutralise it. Therefore, we investigated if this pathway can control other pathogens that can persist intracellularly. When bone marrow-derived macrophages (BMDMs) from wild type, Rab32 or BLOC-3-deficient mice were infected with *Staphylococcus aureus* (*S. aureus*), we observed a significant increased survival in BMDMs deficient for the HPS4 protein, one of the two subunits of BLOC-3, or Rab32 when compared to wild type BMDMs (Fig. 1a and Fig. S1). In line with this, Rab32 is recruited to the vacuole containing *S. aureus* in wild-type but not HPS4-deficient BMDMs (Fig 1B). Given that *S. aureus* is a Gram-positive bacterium, these data suggested that Rab32 and BLOC-3 are components of a broad antimicrobial pathway. Therefore, we investigated if this pathway can also limit infection by fungal pathogens. Wild-type and HPS4-deficient mice were infected with *Candida albicans*, and the fungal burden in kidneys, the main organ affected by this pathogen, was evaluated 72 hours post-infection. BLOC-3-deficient mice exhibited a 14-fold increase in kidney fungal CFUs (Fig 1C). These results indicate that the Rab32/BLOC-3-dependent pathway is critical to defend the host from both bacterial and fungal attacks.

**Fig 1.**
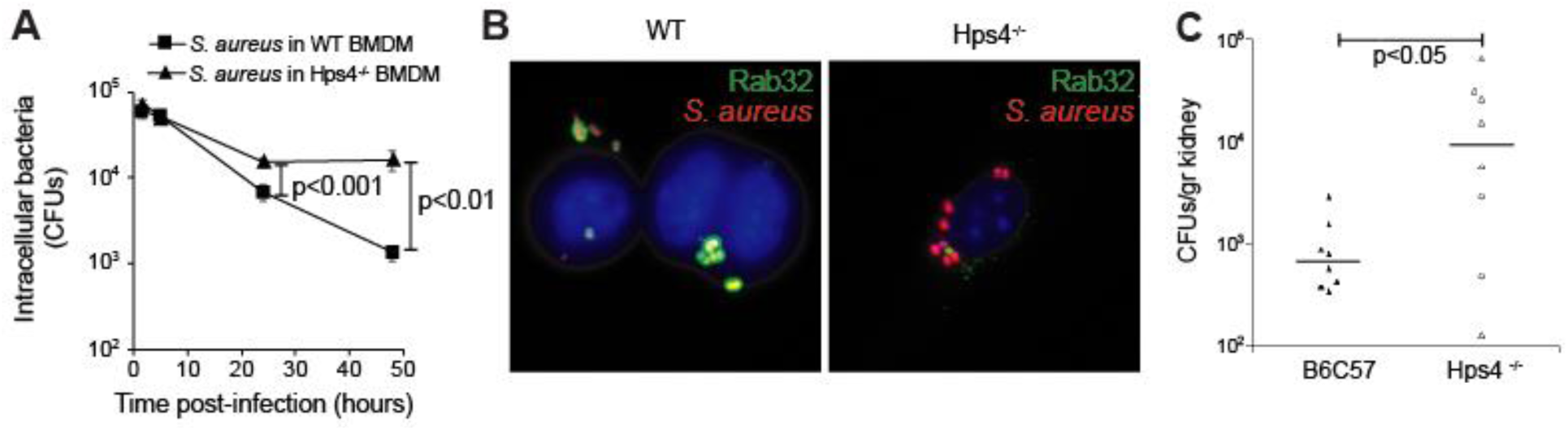
The The Rab32/BLOC-3 pathway mediates the killing of different pathogens. (A) and (B) Mouse bone marrow derived macrophages (BMDM) were derived from control mice C57BL/6 (wt) or from HPS4^-/-^ mice, infected with *S. aureus* and (A) CFUs were enumerated at the times indicated or (B) cell were fixed at 3 hours post infection (hpi) and stained to show Rab32 localisation. (C) wt or HPS4^-/-^ mice were infected with *Candida albicans*, and fungal burden in kidneys was evaluated 72 hours post-infection.

To investigate potential mechanisms for the Rab32-dependent clearance, we infected BMDMs from wild-type and mice defective in particular antimicrobial factors with *S*. Typhi. ROS are molecules that are toxic to all species that have not evolved strategies to neutralise them(Imlay, 2008) Innate immune cells can assemble phagocytic NADPH oxidase on the phagosome to generate ROS to kill intracellular pathogens (Rhen, 2019). BMDMs derived from NADPH oxidase-deficient mice (Phox-/-) clear *S*. Typhi similarly to wild-type BMDM, while a *S*. Typhi strain engineered to deliver the protease GtgE that cleaves Rab32 (Spanò and Galan, 2012) is not killed so efficiently in either macrophages (Fig. 2A). Similarly, the production of iNOS and the cathelicidin-related antimicrobial peptide (Cramp), two important mechanisms that control pathogenic species, are not essential to clear *S*. Typhi in murine BMDMs (Fig. 2B, C). These data indicate that the Rab32 pathway does not require these well characterized mechanisms of pathogen clearance and that another novel mechanism underpins that ability of the Rab32 pathway to clear *S*. Typhi infections in murine cells.

**Fig 2.**
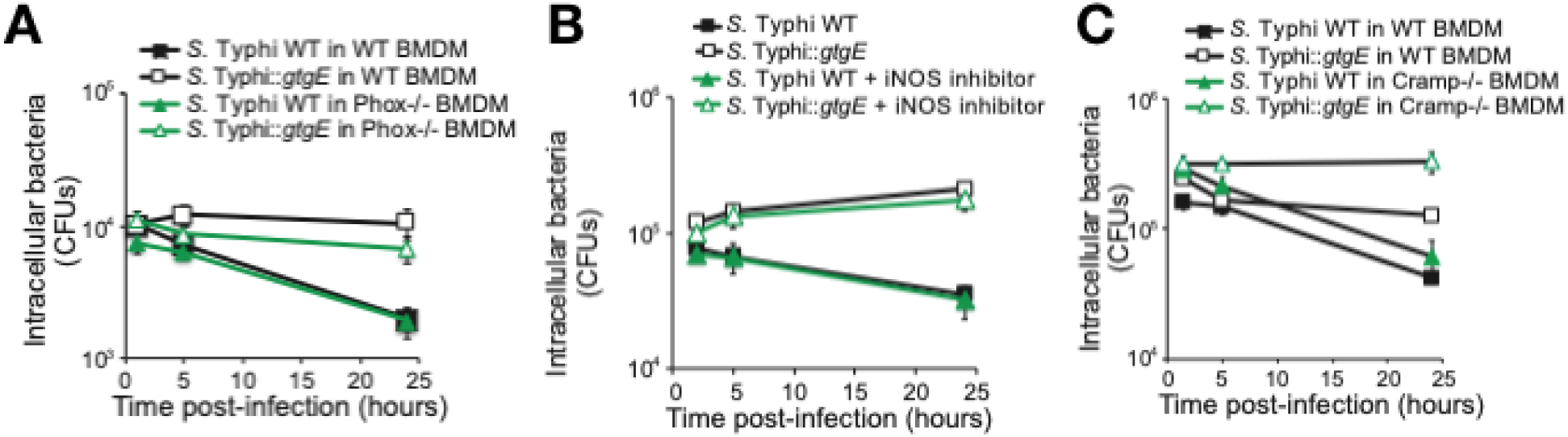
The The Rab32/BLOC-3 pathway do not require oxidative burst to clear bacterial and fungal infections in murine cells. BMDM were infected with S. Typhi wild-type (WT) or expressing GtgE (*::gtgE*) and CFUs were enumerated at the times indicated. (A) BMDM were derived from control mice (wt) or from NADPH oxidase ^-/-^ mice (Phox-/-). (B) BMDM were infected in the presence or absence of the iNOS inhibitor 1400W. (C) BMDM were derived from control mice (wt) or CRAMP^-/-^ mice.

The broad-host *Salmonella* serovar *S*. Typhimurium delivers two type III secretion effectors GtgE and SopD2 that confer the ability of isolates of this serovar to infect mice (Spanò and Galan, 2012; Spanò, 2016; Solano-Collado et al., 2018). *S*. Typhi lacks these two effectors and cannot infect mice but is able to survive in human macrophages and cause a systemic infection in humans. These facts could suggest that the Rab32/BLOC-3 host-defense pathway is not fully active in human macrophages. However, the Rab32 and BLOC-3 genes are present in humans and, genome-wide association studies have shown that single nucleotide polymorphisms in the Rab32 untranslated regions are associated with increased susceptibility to leprosy, a human bacterial infection caused by the intracellular bacterium *Mycobacterium leprae* (Zhang et al., 2009; Liu et al., 2015), suggesting that Rab32 could be part of a pathway critical to control some bacterial infection in humans. Two scenarios could explain these findings: a) Rab32/BLOC-3 are not part of a host-defence pathway in humans or b) a Rab32/BLOC-3 pathway is active in humans as an anti-microbial pathway but *S*. Typhi has evolved molecular strategies to evade it.

To assess if the Rab32/BLOC-3 host-defence pathway is active as an antimicrobial pathway in humans, we investigated the requirement of Rab32 and BLOC-3 in controlling bacterial growth in human macrophages. We used a *S*. Typhi strain engineered to express the *S*. Typhimurium type III secretion effector GtgE, a specific protease that cleaves the three Rab GTPases, Rab32, Rab29 and Rab38 (Spanò and Galan, 2012; Spanò et al., 2011). We infected human macrophage-like THP-1 cells with a *S*. Typhi wild-type isolate or an isogenic strain engineered to express GtgE (*S*. Typhi::*gtgE*). GtgE delivery from *S*. Typhi results in the cleavage of human Rab32 (Fig. 3A), indicating that GtgE can target endogenous human Rab32, in agreement with the previous observation that GtgE cleaves ectopically expressed human Rab32 (Spanò and Galan, 2012). When we infected human blood-monocyte derived primary macrophages, we observed that Rab32 localises on the surface of the vacuoles containing wild-type *S*. Typhi, but is mostly absent from the surface of the vacuoles containing *S*. Typhi::*gtgE* (Fig 3B,C).

**Fig 3.**
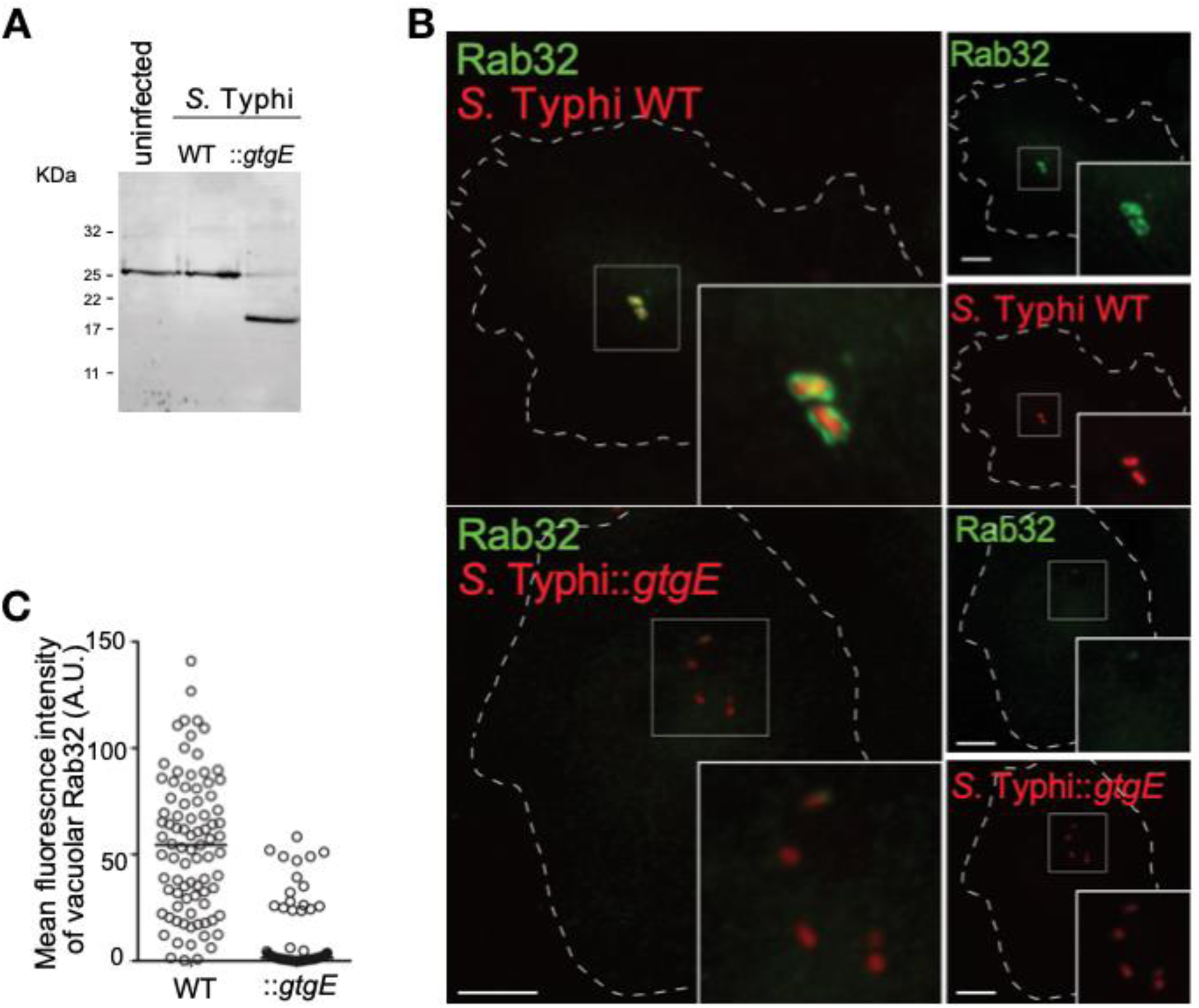
GtgE delivery from *S*. Typhi results in the cleavage of human Rab32. (A) PMA-differentiated THP-1 cells were left uninfected or infected with either wild-type *S*. Typhi (WT) or a *S*. Typhi strain expressing GtgE (::*gtgE*). Cells were lysed 2.5 hours post-infection (p.i.) and analyzed by western blot with a Rab32 specific antibody. (B) and (C) Peripheral blood monocyte-derived macrophages were infected with either wild-type *S*. Typhi (WT) or a *S*. Typhi strain expressing GtgE (::*gtgE*), both carrying a chromosomal copy of the *mCherry* gene, fixed at 2.5 hours p.i. and analyzed by immunofluorescence with a Rab32 specific antibody. Scale bar=10 µm.

We then investigated if this removal of Rab32 from the bacterial vacuole has any effect on *S*. Typhi survival. GtgE expression confers *S*. Typhi a 3-fold replicative advantage in blood-monocyte derived primary macrophages at 24 hours post-infection (Fig 4A), suggesting that one of the three Rab GTPase targeted by GtgE (Spanò et al., 2011; Spanò and Galan, 2012) controls *S*. Typhi intracellular survival in human macrophages. As Rab38 mRNA is hardly detectable in either THP-1 or primary macrophages (data not shown and “The Human Protein Atlas”), we analyzed if either Rab32 or Rab29 are responsible for the limitation of *S*. Typhi growth in human macrophages by knocking down either Rab29 or Rab32 from THP-1 cells (>70% and 75% knockdown respectively; data not shown). While depletion of Rab32 resulted in a significantly increased replication of *S*. Typhi (Fig 4B), a slightly reduced replication was instead observed when Rab29 was depleted. Taken together, these results indicate that Rab32 is critical to control *S*. Typhi infections in human macrophages.

**Fig 4.**
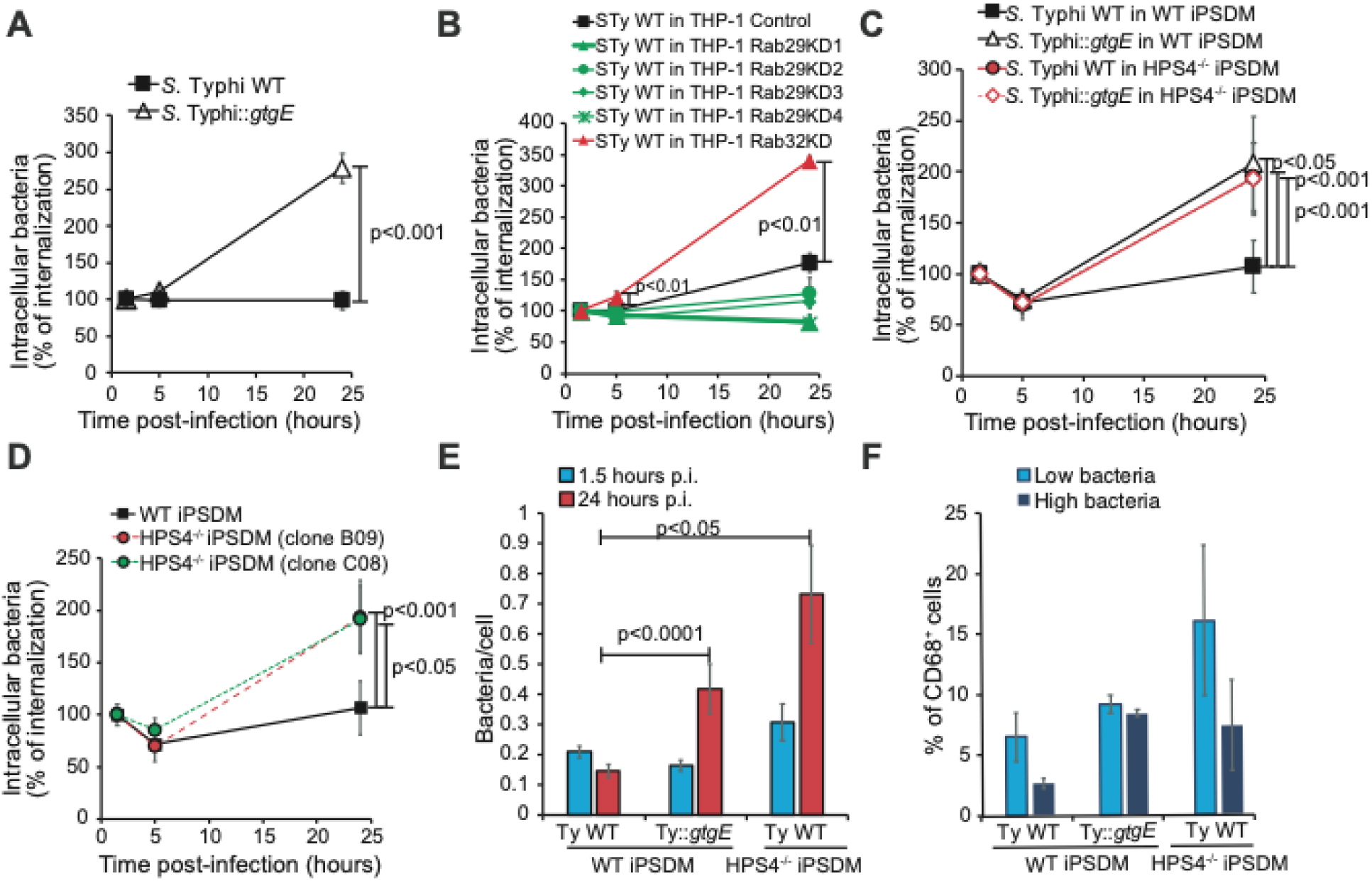
Rab32 inactivation and BLOC-3 knock-out results in *S*. Typhi over-replication in human macrophages. (A) Peripheral blood monocyte-derived macrophages were infected with either wild-type *S*. Typhi (WT) or a *S*. Typhi strain expressing GtgE (::*gtgE*). Cells were lysed at the indicated time points to measure CFUs. (B) THP-1 cells were transduced with lentivirus to silence the indicated Rab, infected with wild type *S*. Typhi and lysed at the indicated time points to measure CFUs. (C) Human macrophages derived from WT or HPS4^-/-^ hiPSCs, were infected with either wild-type *S*. Typhi (WT) or a *S*. Typhi strain expressing GtgE (::*gtgE*), lysed at the indicated time points for counting of intracellular CFUs. (D) Human macrophages derived from WT or HPS4^-/-^ hiPSCs were infected with wild-type *S*. Typhi (WT) and lysed at the indicated time points to measure CFUs. (E) Human macrophages derived from WT or HPS4^-/-^ hiPSCs were plated on glass coverslips, then infected with *S*. Typhi *glmS∷Cm::mCherry* or *S*. Typhi::*gtgE glmS∷Cm::mCherry* and fixed at 1.5 and 24 hours p.i. Differentiated macrophages were identified by CD68 staining, bacteria in CD68-positive cells were counted and the average value plotted. At least 120 cells were counted for each timepoint in two independent experiments. (F) Human macrophages derived from WT or HPS4^-/-^ hiPSCs, were infected with *S*. Typhi *glmS∷Cm::mCherry* or *S*. Typhi::*gtgE glmS∷Cm::mCherry*, fixed at 24 hours p.i. and analyzed by flow cytometry. CD68+ cells were sub-gated in two population containing respectively low or high mCherry signal (i.e. bacterial content). Results in (A), (B), (C) and (D) are reported as percentage of CFUs measured at the first time point (1.5 hours p.i.). Values are means ± SEM of at least three independent experiments performed in triplicates. p-values were calculated using the Student’s t-test and are indicated only when <0.05

We then used macrophages derived from human-inducible pluripotent stem cells (hiPSCs), a recently established model for the study *Salmonella* infection (Hale et al., 2015; Alasoo et al., 2015). First, we confirmed that, similarly to what observed in THP-1 and primary macrophages, GtgE expression also confers an advantage to *S*. Typhi in hiPSC-derived macrophages (Fig 4C). Next, we used CRISPR/Cas9 technology to generate hiPSCs deficient for HPS4. As shown in Fig 4D, macrophages derived from two independent clones of HPS4-deficient hiPSCs have a significant increased number of *S*. Typhi intracellular CFUs at 24 hours post-infection, demonstrating that BLOC-3 is important to limit *S*. Typhi growth in human macrophages. To obtain further insights into the effects of GtgE expression in *S*. Typhi or the knock-out of BLOC-3 in these macrophages, we measured the number of *S*. Typhi in single cells by immunofluorescence (Fig 4E) or flow cytometry analysis (Fig 4F). These experiments confirmed that removal of either Rab32, obtained through GtgE delivery, or BLOC-3 results in an increased percentage of host cells containing higher numbers of bacteria. This indicates that both Rab32 and its guanine nucleotide exchange factor BLOC-3 are required for the control of *Salmonella* survival and replication in human macrophages. We also observed that although *S*. Typhi has a replicative advantage in the absence of BLOC-3, the expression of GtgE does not confer any significant additional advantage (Fig. 4C), in agreement with the model that Rab32 and BLOC-3 are components of the same pathway. The results of these experiments indicate that the Rab32/BLOC-3 pathway is active as a host-defence pathway in human macrophages and can limit *S*. Typhi replication.

To test if the human Rab32/BLOC-3 pathway exerts broad antimicrobial activity, we infected macrophages derived from two independent clones of HPS4-deficient hiPSCs with *S. aureus*. As shown in Fig. 5A, HPS4 knock out results in ≈ 10-fold increased survival of *S. aureus* in human macrophages. However, in contrast to *S. aureus* (Fig 5A) and other pathogens, such as *E. coli* O157 (Fig. 5B), *S*. Typhi is not as efficiently cleared by wild-type human macrophages during infection, but instead persist in the majority of infected cells (Fig. 5B and Fig. 4A-D). Therefore, we hypothesized that *S*. Typhi actively counteracts the Rab32/BLOC-3 pathway. Since the broad-host *S*. Typhimurium neutralizes this pathway through the action of effectors delivered by type III secretion systems, we tested if *S*. Typhi survival in wild-type human macrophages is dependent on *S*. Typhi type III secretion systems. We observed that *S*. Typhi survival in hiPSC macrophages is dependent on its SPI-1 (Fig 5C), but not on its SPI-2 type III secretion system (Fig S2). Indeed, a SPI-1 type III secretion system mutant of *S*. Typhi (*S*. TyphiΔ*invA*) was unable to survive in macrophages derived from hiPSCs (Fig 5C), in agreement with published results (Forest et al., 2010). Interestingly, *S*. TyphiΔ*invA* survived much better in HPS4-deficient macrophages (Fig 5C), suggesting that the Rab32/BLOC-3 pathway is involved in *S*. Typhi killing and that *S*. Typhi needs this secretion system to counteract this pathway. To confirm that HPS4 removal did not result in a completely impaired bacterial killing, we infected HPS4-deficient macrophages with pathogenic *E. coli* O157. This pathogen is not able to survive in either wild type or HPS4-deficient macrophages (Fig 5D). These results indicate that *S*. Typhi is able to target the human Rab32/BLOC-3 pathway likely through expression of the SPI-1 type III secretion system. As *S*. Typhi cannot neutralise the mouse Rab32/BLOC-3 pathway, we suggest that *S*. Typhi targets a human-specific component of this pathway.

**Fig 5.**
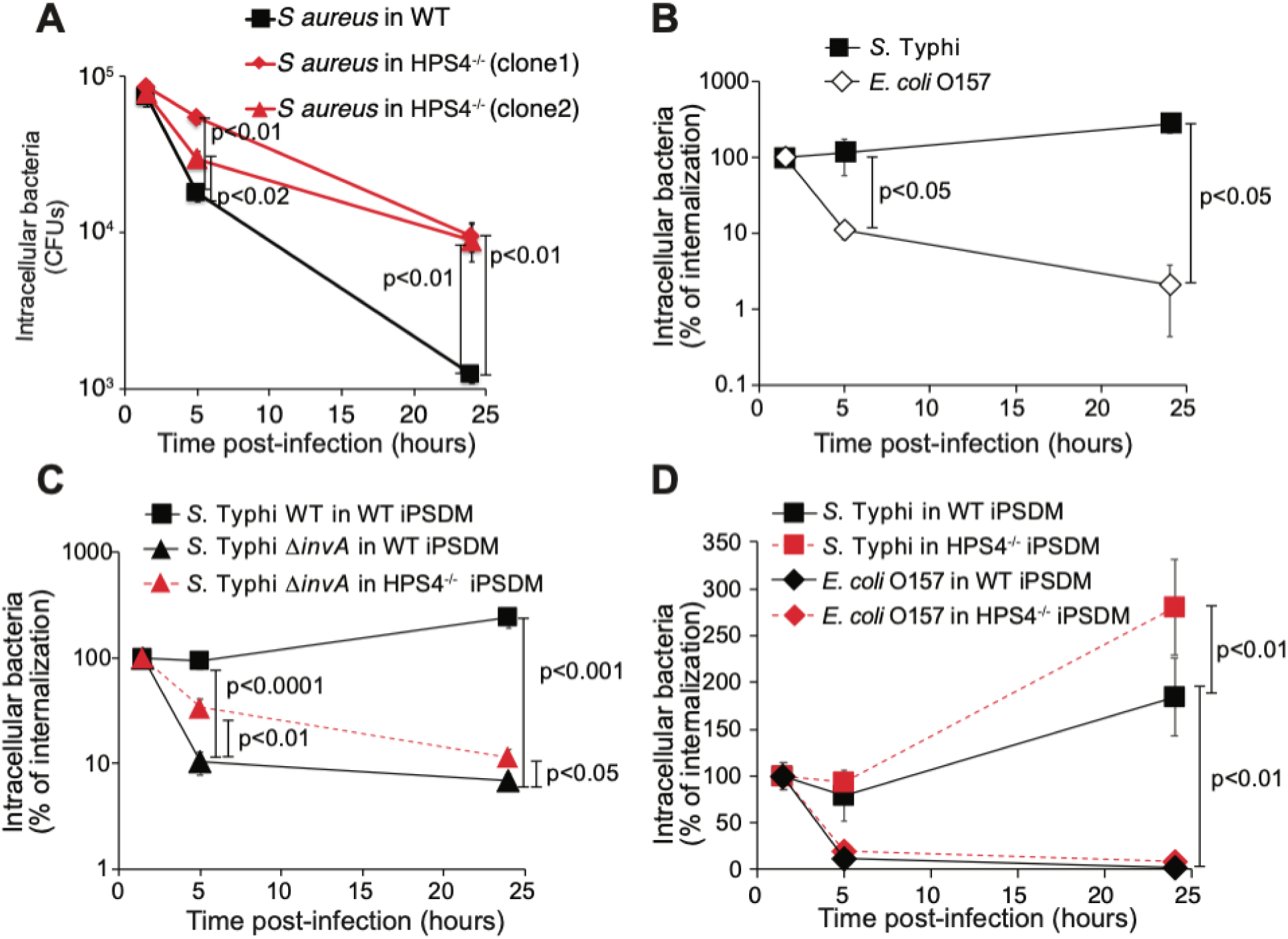
*S*. Typhi counteracts the Rab32/BLOC-3 pathway through the SPI-1 type III secretion system. (A) Human macrophages derived from WT or HPS4^-/-^ hiPSCs were infected with *S. aureus* and lysed at the indicated time points to measure intracellular CFUs. (B) Human macrophages derived from WT hiPSCs were infected wild-type *S*. Typhi (WT) or *E. coli* O157 and lysed at the indicated time points to measure intracellular CFUs. (C) Human macrophages derived from WT or HPS4^-/-^ hiPSCs were infected with either wild-type *S*. Typhi (WT) or *S*. Typhi Δ*invA* (Δ*invA*). Cells were lysed at the indicated time points to measure intracellular CFUs. (D) Human macrophages derived from WT or HPS4^-/-^ hiPSCs were infected with either wild-type *S*. Typhi (WT) or *E. coli* O157. Cells were lysed at the indicated time points to measure intracellular CFUs. Results in (B), (C), and (D) are reported as percentage of CFUs measured at the first time point (1.5 hours p.i.). Values are means ± SEM of at least three independent experiments performed in triplicates. p-values were calculated using the Student’s t-test and are indicated only when <0.05.

## DISCUSSION

Overall, we show that the Rab32/BLOC-3 pathway regulates a broad host defense activity against a variety of intracellular pathogens, including Gram-negative and Gram-positive bacterial pathogens and the fungal pathogen *Candida albicans*. This pathway is critical for the clearance of *S. aureus* and *S*. Typhi in mouse macrophages and for the clearance of *S. aureus* in human macrophages. We show here that this antimicrobial mechanism does not require the production of reactive oxygen species by the phagocytic NADPH oxidase, an ancient and broad antimicrobial mechanism active in macrophages. We also demonstrate it does not require production of nitric oxide by iNOS and the macrophage antimicrobial peptide CRAMP. Remarkably and in contrast to *S. aureus, S*. Typhi uses its SPI-1 type III secretion system to evade this pathway in humans and persist in human macrophages.

Not dissimilar from *S*. Typhimurium, which has evolved to deliver the protease GtgE and the Rab GAP SopD2 that act redundantly to inactivate the murine Rab32/BLOC-3 trafficking pathway, *S*. Typhi appears to have evolved a strategy to target the human Rab32/BLOC-3 pathway that requires its SPI-1 type III secretion system. A possible reason for evolving a different strategy could rely on the fact that GtgE also targets Rab29, which is required for the efficient delivery of typhoid toxin from *S*. Typhi-infected cells (Spanò et al., 2011). Therefore, we speculate that GtgE, although able to cleave the human Rab32, would not confer overall an advantage to *S*. Typhi because it would interfere with other pathogenic features of this bacterium, including the delivery of typhoid toxin. Similar to *S*. Typhimurium, *S*. Typhi may have evolved a number of redundant effectors with different activities to target the Rab32/BLOC-3 pathway.

In conclusion, in this manuscript we reveled as the Rab32/BLOC-3 pathway is a novel, general and broad-spectrum antimicrobial pathway that controls a variety of infectious pathogens and showed that *S*. Typhi use its SPI-1 type III secretion system to counteract this pathway and survive in human macrophages.

**This paper is part of Stefania Spanò’s scientific legacy and this would have not been possible without her intelligence, vision and persistence. A dreadful destiny has snatched her from us too early, but her discoveries and ideas are living and flourishing**.

## MATERIALS AND METHODS

### Bacterial Strains and Plasmids

The wild-type *Salmonella enterica* serovar Typhi strain ISP2825 have been previously described (Galán and Curtiss, 1991). All the *S*. Typhi deletion strains were constructed by standard recombinant DNA and allelic exchange procedures as previously described (Kaniga *et al*., 1994) and are listed in Table S1. All the plasmids used in this study were constructed using standard recombinant DNA techniques and are listed in Table S2. *S*. Typhi *glmS∷Cm::mCherry* and *S*. Typhi::*gtgE glmS::Cm::mCherry* that constitutively express *mCherry* from a single chromosomal copy at the *att*Tn7 site were generated by P22 transduction using phages obtained from the *S*. Typhimurium SL1344 *glmS::Cm::mCherry* (a gift from Leigh Knodler; (Knodler et al., 2014))

**Table S1.** List of *S*. Typhi strains used in this study.

| Strain | Genotype | References |
| --- | --- | --- |
| ISP2825 | wild type | Galán and Curtiss, 1991 |
| SB2522 | ::*gtgE* | Spanò and Galán, 2012 |
| SB2174 | $\Delta invA$ | Spanò *et al.*, 2011 |
| SB1958 | $\Delta spiA$ | Spanò *et al.*, 2011 |
| SBB001 | *glmS::Cm::mCherry* | this study |
| SBB002 | ::*gtgE glmS::Cm::mCherry* | this study |

**Table S2.** List of plasmids used in this study.

| Plasmid | Description | References |
| --- | --- | --- |
| pVSVG | pVSV-G | Spanò *et al.*, 2011 |
| pGag/Pol | pMLV-Gag-Pol | Spanò *et al.*, 2011 |

### Cell culture

THP-1 cells were maintained in RPMI 1640 medium (Invitrogen), 10% Fetal Bovine Serum (FBS, Invitrogen), 2 mM Glutamine (Invitrogen), 1 mM Sodium Pyruvate (Invitrogen) and 10 mM Hepes (Invitrogen). The cells were maintained at a concentration between 0.1 and 1 million cell/ml. THP-1 differentiation was induced by adding 100 nM Phorbol 12-Myristate 13-Acetate (PMA) for or 48 hours before infection. When indicated, THP-1 differentiated cells were treated with human-INFγ before infection. HEK293T cells were maintained in DMEM high glucose 2 mM glutamax (Invitrogen), 10% FBS.

Blood was collected from healthy human volunteers, according to procedures approved by the Life Science and Medicine College Ethics Review Board of the University of Aberdeen (CERB/2016/11/1299). Peripheral blood monocyte-derived macrophages were prepared as described in (Arnold et al., 2014) with some modifications. Briefly 13 ml of blood were collected from each donor, diluted to 35 ml with Hank’s balance salt solution (HBSS, Invitrogen), then loaded onto 15 ml of Lymphoprep™ (Stem Cell Technology) for the separation of the peripheral blood mononuclear cells. Isolated peripheral blood mononuclear cells were resuspended in DMEM containing 10% autologous human serum (freshly prepared from the same donor) and seeded on coverslips or tissue-treated plastic. Cells were plated at 5×10^5^/well in 24-well plates. After 24 hours the non-adherent cells were removed, fresh medium added, and the cells left for 7-9 days to differentiate.

Undifferentiated human induced pluripotent stem cells line (KOLF2-C1) was maintained on a monolayer of mitotically inactivated mouse embryonic feeder (MEF) cells in advanced Dulbecco’s modified Eagles/F12 medium (DMEM/F12), supplemented with 20% knockout replacement serum (KSR, Invitrogen), 2 mM L-Glutamine, 0.055 mM β-mercaptoethanol (Sigma-Aldrich) and 8 ng/ml recombinant human FGF2 (RnD Systems), as described previously (Hale et al., 2015). These cells were differentiated into macrophages as described in a previously published method (Hale et al., 2015).

### Candida albicans infection in wild-type or HPS4^-/-^ mice

*<u>Candida albicans</u>* <u>(strain SC5314) was serially grown overnight at 30 °C with shaking. Yeast cells were washed in</u> phosphate buffer saline (PBS, Sigma-Aldrich)<u>, counted and injected intravenously via the lateral tail vein. Animals were infected with 2× 105c.f.u. For analysis of fungal burdens in the kidneys, animals were euthanized 72 hours post infection. Kidneys weighed, homogenized in PBS and serially diluted before plating on to YPD agar supplemented with penicillin/streptomycin (Invitrogen). Colonies were counted after incubation at 37 °C for 24–48 h.</u>

### CRISPR/Cas9 targeting of HPS4

Isogenic intermediate targeting vectors for HPS4 were generated using isogenic and haplotype specific DNA by PCR amplification of KOLF2-C1 gDNA. Firstly, a PCR fragment including homology arms and the critical exon of HPS4) was amplified from KOLF2-C1 gDNA using the following primers; f5F <u>gccagtgaattcgat</u>atacctgccttcttgaactgttttg and f3R <u>tacgccaagcttgat</u>ttaaattgtgctctgtgtgttcctc. The first 15nt of each primer (underlined) served to mediate fusion with the intermediate targeting vector backbone, puc19_RV using the In-Fusion HD Cloning Plus kit (TAKARA). The HSP4 amplicon was purified and 75ng incubated with 50ng of EcoRV digested puc19_RV vector for 15 minutes at 50°C and transformed into Stellar competent cells (TAKARA). Positive clones were verified by Sanger sequencing. To replace the critical exon with the gateway R1-*pheS/zeo*-R2 cassette, sequence verified clones were electroporated with the pBABgbaA plasmid (Tate and Skarnes, 2011). This was then maintained in tetracycline (5 μg/ml) at 30°C. Early log phase cultures were induced to express the red operon following addition of 0.1% arabinose and incubation for 40 min at 37°C. From these cultures electro competent cells were prepared as previously described(Tate and Skarnes, 2011). The R1-*pheS/zeo*-R2 cassette was amplified using the following primers; U5 <u>ttagtggtgtcagcagttctgagtatagagaggtagaatagtcccaagcc</u>aaggcgcataacgataccac and D3 <u>agttgtgcagcaagggaatggggctggaagaaaggggtctggagttactc</u>ccgcctactgcgactataga. Underlined sequences in each of these primers denotes 50nt of homology towards a region 5’ (U5) or 3’(D3) of the critical exon. This amplicon was purified and 300ng electroporated into the recombination ready verified clones from the first step before selection in carbenicillin (50µg/ml) and zeocin (10 µg/ml). Positive clones were verified by Sanger sequencing. To generate the donor plasmid for precise gene targeting via homology directed repair, the intermediate targeting vectors were turned into donor plasmids via a Gateway exchange reaction. LR Clonase II Plus enzyme mix (Invitrogen) was used to perform a two-way reaction exchanging only the R1-*pheSzeo*-R2 cassette with the pL1-EF1αPuro-L2 cassette as previously described (Tate and Skarnes, 2011). The latter had been generated by cloning synthetic DNA fragments of the EF1α promoter and puromycin resistance cassette into a pL1/L2 vector(Tate and Skarnes, 2011)

As part of the primer design process, two separate guide RNAs (gRNAs) targeting within the same critical exon were selected. The gRNAs were identified using the WGE CRISPR tool (Hodgkins et al., 2015) and were selected based on their off-target scores to minimise potential off target damage. gRNAs were suitably positioned to ensure DNA cleavage within the exonic region, excluding any sequence within the homology arms of the targeting vector. Plasmids carrying single guide (sg) RNA sequences were generated by cloning forward and reverse strand oligos into the BsaI site of either U6_BsaI_gRNA or p1260_T7_BsaI_gRNA vectors (kindly provided by Sebastian Gerety, unpublished). The CRISPR sequences as follows (PAM sequence); left CRISPR (CCA) GCGAGAATGTGAGGGCGAGCG and right CRISPR (CCT) TCAGCAACAACAGGGGCTCC (WGE IDs: 1181940311 and 1181940319 respectively).

To deliver plasmids expressing gRNA, donor templates and Cas9, 2 × 10^6^ KOLF2-C1 cells were nucleofected (AMAXA nucleofector 2B) with 2 µg of donor plasmid, 4 µg hCas9 D10A (Addgene plasmid #41816) (Mali et al., 2013) and 3 µg of gRNA plasmid DNA. Following nucleofection, cells were selected for up to 11 days with 0.25 μg ml^-1^ puromycin. Individual colonies were picked into 96-well plates, expanded and genotyped. Positive insertion of the cassette into the correct locus was confirmed by PCR using cassette-specific primers ER (gcgatctctgggttctacgttagtg) and PNFLR (catgtctggatccgggggtaccgcgtcgag). To determine the presence of deleterious insertions or deletions (indels) around the CRISPR target site of the opposite allele, a PCR amplicon was generated using the primers PR (actagttctaacagctggtggatac) and PF (ttttgcagactgacaactattccag), purified and Sanger sequenced using SR1 (cttctggacaggcctccttg) and SF1 (atatttgccgaaccagccca). To minimize the potential for off-target effects, two independently derived clones, B09 and C08, with specific deletions of 47 and 29 base pairs, respectively, were isolated and used in this study.

### Intracellular growth experiments

Overnight cultures of the different *S*. Typhi strains or *S aureus* (strain SH1000 (Horsburgh et al., 2002)) were diluted 1/20 in LB broth containing 0.3 M NaCl and grown for 2 hours and 45 minutes at 37°C. Cells were infected with the different strains of *S*. Typhi in HBSS at the desired multiplicity of infections. One-hour p.i. cells were washed three times with HBSS and incubated in growth medium supplemented with 100 μg/ml gentamicin for 30 min to kill extracellular bacteria. Cells were then washed with HBSS, and fresh DMEM containing 5 μg/ml gentamicin was added to avoid cycles of reinfection. At the indicated time points the cells were washed twice in PBS and the intracellular bacteria recovered lysing the cells in 0.1% sodium deoxycholate (*Salmonella*) or 0.1% Triton X-100) (Sigma-Aldrich) (*S. aureus*) in PBS and counted by plating serial dilutions on LB-agar plates.

### Western blot

PMA-differentiated THP-1 cells were infected as described above and lysed in SDS-PAGE loading buffer 2.5 hours post-infection. Western blot analysis was performed using Odyssey infrared imaging system (LI-COR Biosciences). The following antibodies were used for Western blot analysis: rabbit polyclonal anti-Rab32 (GeneTex, 1:1,000 dilution); Donkey anti-rabbit IR Dye 800 (Li-COR, 1:10,000 dilution).

### Rab32 and Rab29 knockdown in THP-1

THP-1 were transduced with lentivirus expressing shRNA targeting Rab29 (TRCN0000299449; TRCN0000303685; TRCN0000381042; TRCN0000303621 Sigma-Aldrich) or Rab32. (TRCN0000047746 Sigma-Aldrich). Twenty-four hours post-transduction, cells were treated with 5 µg/ml puromycin to kill of the non-transduced cells and kept in culture for not more than two weeks. 72 hours before the infection the cells were treated 100 nM PMA for 48 hours to induce differentiation. 24 hours before the infection the PMA was removed and the cells stimulated with 100 ng/ml of human interferon gamma (INFγ) and finally then infected with *S*. Typhi as described above.

### Immunofluorescence

Bone marrow derived mouse macrophages, human monocyte-derived macrophages and wild-type or HPS4 deficient iPSDM were plated on glass coverslips (#1 Thermo) infected with different *S*. Typhi strains or with *S aureus* (strain SH1000 (Horsburgh et al., 2002)) and fixed at the indicated times p.i. with 4% paraformaldehyde (PFA) for 10 minutes. Cells were then permeabilized for 20 minutes by incubating in 0.02% Saponin (Sigma-Aldrich), 0.2% bovine serum albumin (BSA, Sigma-Aldrich), 50 mM NH_4_Cl (Sigma-Aldrich) in PBS and incubated for 1 hour with monoclonal mouse anti CD68 (KP1 Invitrogen, 1:200 dilution). Alternatively, cells were permeabilized for 20 minutes by incubating in 0.2% Triton X-100 (Sigma-Aldrich), 0.2% bovine serum albumin (BSA, Sigma-Aldrich), 50 mM NH_4_Cl (Sigma-Aldrich) in PBS and incubated for 1 hour with a rabbit polyclonal anti-Rab32 (GeneTex, 1:200 dilution). Cells were then stained using the appropriate Alexa Fluor^®^ 488 or Alexa Fluor^®^ 555-conjugated secondary antibodies (Invitrogen). Images were acquired either using in a Nikon (Eclypse Ti2) equipped with a CFI Plan Apochromat 100x objective and a Prime 95B 25 mm CMOS camera (Photometrics) or a PerkinElmer Spinning disk confocal equipped with an ORCA Flash 4.0 CMOS camera (Hamamatsu). Images were analyzed using the respective software (Nikon Elements or Volocity).

### Flow cytometry

hiPSC-derived macrophages wild type or HPS4-deficient were plated on non-tissue culture treated 6-well plates (Thermo Fisher Scientific) and infected with *S*. Typhi *glmS::Cm::mCherry* or *S*. Typhi::*gtgE glmS::Cm::mCherry*. At the indicated time p.i. the cells were detached using 500 µl of Versene (Invitrogen) and mixed with an equal volume of 4% PFA for 5 minutes. Fixed cells were then centrifuged and resuspended in 4% PFA for 5 minutes. The cells were then transferred in flow cytometry tubes, permeabilized 15 minutes in PMZ-S, then incubated for 1 hour with anti-CD68 (1:200) and then with anti-mouse Alexa Fluor^®^ 488. The samples were analyzed by flow cytometry (Fortessa, BD Biosciences) and FlowJo software.

## ACKNOWLEDGEMENTS

We are very grateful to Leigh Knodler for her generous gift of P22 phages from a *S*. Typhimurium *glmS::Cm::mCherry* strain. We would also like to thank Bill Skarnes, Alex Alderton, Vivek Iyer, Mark Thomson, Tristan Thwaites, Oliver Dovey for their help with hiPSC and CRISPR/Cas9 knock-out. This work was supported by the Wellcome Trust (Seed Award 109680/Z/15/Z), the European Union’s Horizon 2020 ERC consolidator award (2016-726152-TYPHI), the BBSRC (BB/N017854/1), the Royal Society (RG150386), and Tenovus Scotland (G14/19) to SS. VSC is recipient of a European Union’s Horizon 2020 research and innovation programme Marie Sklodowska-Curie Fellowship (706040_KILLINGTYPHI). GDB, DMR and IMD were supported by the University of Aberdeen, Wellcome Trust (102705) and the MRC Centre for Medical Mycology (MR/N006364) (currently at the University of Exeter).

