## supplementary material (S1 and S2) for "The Rab32/BLOC-3 dependent pathway mediates host-defence against different pathogens in human macrophages"

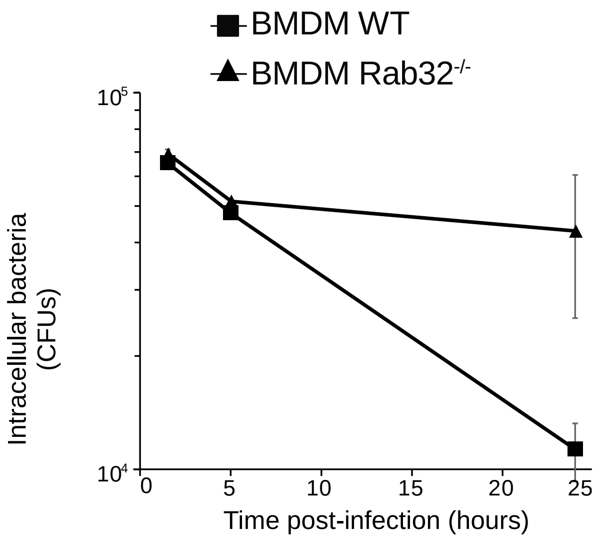


**Fig S1. The Rab32/BLOC-3 pathway mediates the killing of different pathogens.** Mouse bone marrow derived macrophages (BMDM) were derived from control mice C57BL/6 (wt) or from Rab32-/- mice were infected with *S. aureus*. Cells were lysed at the indicated time points to measure intracellular CFUs.

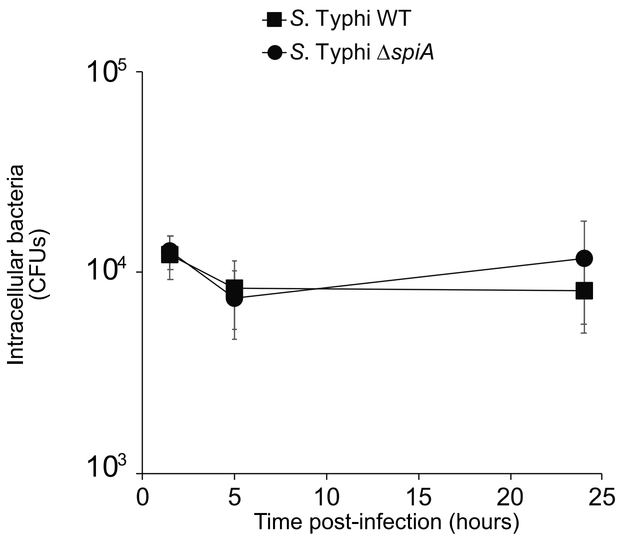


**Fig S2. *S.* Typhi survival in macrophages does not depend on its SPI-2 type III secretion system**. Human macrophages derived from induced pluripotent stem cells were infected with either wild-type *S.* Typhi or *S.* Typhi Δ*spiA*. Cells were lysed at the indicated time points to measure intracellular CFUs.
